# Bridging macroscopic and microscopic modeling of electric field by brain stimulation

**DOI:** 10.1101/2025.04.07.647453

**Authors:** Boshuo Wang, Aman S. Aberra

## Abstract

**Background:** Modeling the electric field (E-field) on the microscopic scale improves our understanding of brain stimulation modalities and modeling methods but requires careful consideration of the conductivity values and correction of the field amplitudes to match conventional models on the macroscopic scale.

**Objective:** We analyze the correction step and discuss its relevance and implications for E-field modeling efforts bridging the macroscopic and microscopic scales.

**Methods:** We provide the theoretical framework for comparing microscopic and macroscopic models and describe approaches for effectively and efficiently matching the E-field amplitude and tissue conductivity.

**Results:** Consistent results can be obtained for brain stimulation models on different scales with appropriately selected conductivity and E-field amplitudes.

**Conclusion:** Microscopic E-field models enable numerical estimation of conductivity of macroscopic homogenous neural tissue from microscopically realistic brain samples and exploration of the effect of microscopic E-field perturbations on neural activation threshold by brain stimulation, therefore improving modeling accuracy.

## Introduction

Recently, Qi et al. developed a computational framework to model electrical stimulation at microscopic levels in large brain tissue samples spanning hundreds of μm. Using the IARPA Phase I scanning electron microscopy dataset in mouse primary visual cortex, they modeled individual neurons and other cell types with realistic morphologies and sub-μm features [1]. The framework enables simulations of the response of a large neuron population to electric fields (E-fields) imposed by electromagnetic stimulation. Crucially, it captures interactions between the imposed E-field and the membranes of the interwoven cellular processes, both within a neuron itself and between neighboring cells. The interaction between dense cellular structures and the imposed E-field generates microscopic perturbations to the E-field, or an E-field “noise” with zero spatial mean. But the magnitude of this noise and its net impact on neural activation is unknown. In contrast, the vast majority of computational studies of brain stimulation, including our own [2], used a “two-step approach” which 1) solved the E-field distribution at the macroscopic level, assuming negligible microscopic E-field noise, and then 2) coupled the E-field distribution to compartmental neuron models as an extracellular potential. Qi et al. explored how the E-field noise affects neural activation thresholds and examined the validity of the long-established two-step approach to simulate the neural response [3]. Their work presents an impressive effort to improve our understanding of brain stimulation and modeling methodology, and it provides the first major and detailed computational validation of the conventional two-step approach that has been used ubiquitously in the field of neuromodulation.

In the study, the authors imposed uniform E-fields across the block of tissue (250×140×90 µm^3^) and compared activation thresholds of individual neurons subjected to 1) the total E-field including perturbations from the surrounding membranes or 2) a uniform E-field modeled in isolation in a homogenous extracellular space, without E-field noise. Although E-field perturbations initially reduced the thresholds significantly by 15% to 45% (30% on average), proper correction for the imposed E-field amplitude and matching of tissue conductivity (section 2.8 of [1]) resulted in only minimal changes (less than 10%), therefore mostly supporting the conventional two-step approach. Here, we further analyze and discuss this correction step and its relevance to E-field modeling efforts bridging the macroscopic and microscopic scales, summarize more efficient alternative methods in the literature, and outline future applications of this work.

### Theoretical framework

In the two-step approach, the first step utilizes macroscopic finite or boundary element methods (FEM/BEM) to solve the E-field in the brain tissue, which is modeled using homogenous tissue volumes with conductivity representing the “bulk average” of the underlying cellular composition (Fig. 1A). The conductivity is obtained from experimental measurements [4] and for neural tissue like grey matter (GM) is typically in the range of 0.1–0.3 S/m [5,6]. Inhomogeneity of tissue conductivities within the same tissue type, if considered, would be on the macroscopic scale and gradual, and therefore different from microscopic conductivity contrast, e.g., between the extracellular space and membranes. The macroscopic E-field solution is sampled on the microscopic scale from locations where the neuron models represented by one dimensional cables are virtually “re-inserted” in the tissue (Fig. 1B, dotted lines), and is then transferred to the second step to simulate the neural response [2], e.g., using the NEURON solver.

**Figure 1.**
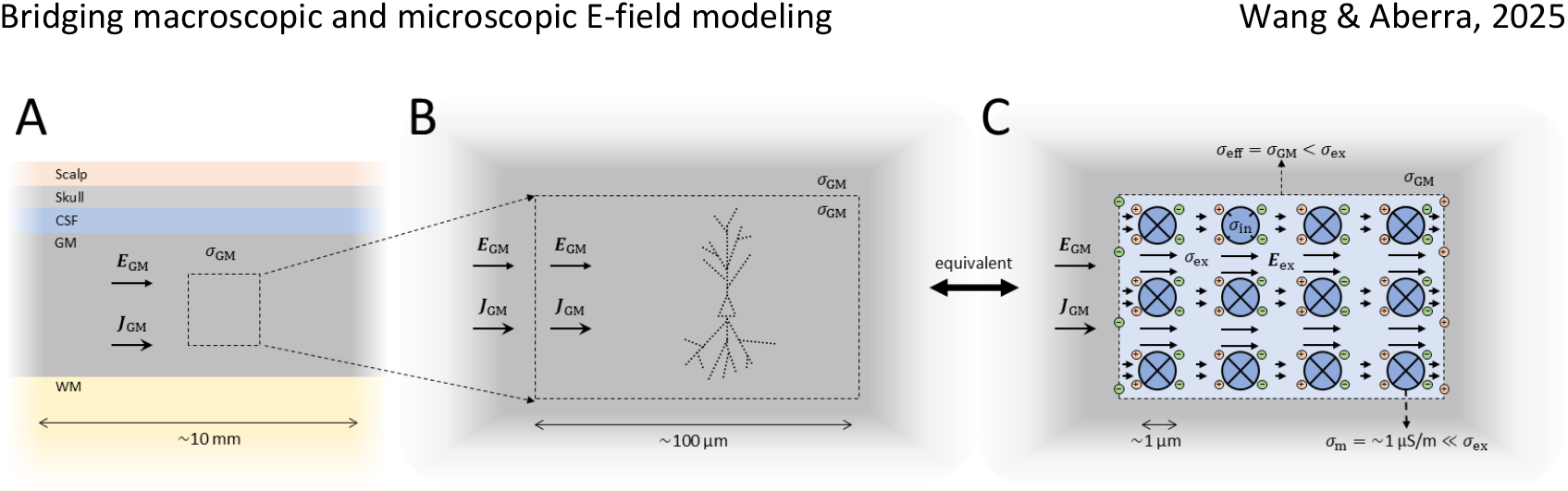
**A**. The first step of the conventional two-step modeling approach utilizes homogenous tissue volumes with bulk tissue conductivity *σ* to simulate the E-field ***E*** and current density ***J*** in the tissue. Five tissue compartments are shown for a head model, with neural tissue like grey matter (GM) having conductivity typically in the range of 0.1–0.3 S/m. CSF: cerebrospinal fluid. WM: white matter. **B**. For the neural simulation in the second step, the E-field or potential is sampled from a region of interest (tissue block shown with dashed outline) on the meso- or microscopic scale where the neuron (dotted outline) is virtually placed. Therefore, the coupling of the E-field to the neuron model is unidirectional, with the influence of the cell membrane on the E-field itself ignored. **C**. The same block of tissue with its underlying cellular structure shown, here represented with circles. The extracellular space (light blue) and intracellular space (dark blue) consist of physiological saline and have higher conductivities *σ*_ex_ and *σ*_in_ (1–2 S/m, e.g., 1.6–1.8 S/m for CSF [5]) compared to bulk neural tissue, whereas the membrane has very low conductivity *σ*_m_ (0.1–1 μS/m, ∼5 nm thickness) and essentially makes the intracellular space inaccessible to current flow (as indicated by the crosses) after initial polarization [7]. Current continuity under quasi-static conditions [8] results in polarization charges on the cell membranes and on the boundaries between the extracellular space and the bulk tissue. These charges (red for positive and green for negative) reduce the average E-field strength in the extracellular domain inside the microscopic tissue block compared to that in the corresponding homogeneous tissue and the surrounding homogenous environment. Therefore, matching the total current through the tissue block and the boundary conditions for the potentials between the two situations are matched, resulting in the same effective conductivity *σ*_eff_ as the bulk tissue. Panel C is adapted from [1] (Figure 2 of the bioRxiv preprint version 5).

The homogeneous tissue consists of microscopic cellular structures embedded in extracellular fluids (Fig. 1C). To be theoretically consistent, replacing the homogenous tissue with the microscopic sample in a macroscopic model (e.g., full head) should not change the E-field solution in the surrounding tissue, except for microscopic noise near the boundary that is negligible on the macroscopic scale. Compared to bulk neural tissue, extracellular fluids have much higher conductivity values in the range of 1–2 S/m, e.g., 1.6–1.8 S/m for cerebrospinal fluid (CSF) [5,9,10]. The high conductivity contrast between the CSF and surrounding bulk tissue, with a ratio from 3 to over 10, results in polarization charges (Fig. 1C, in green and red) on the boundary. Similarly, polarization charges also build up on the neural membranes [1] due to their high conductivity contrast with the extracellular space, reaching a state called “initial polarization” within ∼1 µs [7,11]. Qi et al.’s numerical method calculated charge buildup at the end of this initial polarization phase, which allowed them to treat membranes as nonconducting and provided the initial condition for subsequent simulations of membrane dynamics (i.e., ionic current flow and voltage changes). Together, the charges on the tissue boundary and cell membranes generate a secondary E-field which opposes the imposed (primary) E-field and results in an overall weaker total E-field in the extracellular space compared to that in the equivalent homogeneous tissue. Thus, applying an E-field to a tissue model with microscopic structure results in the same current flow and potential drop as in the homogenous tissue but with lower average E-field and higher current density. The higher current density arises as the currents flow along longer, meandering paths (typically between 1.2 to 2 longer for brain tissue [12]) through the small volume of high conductivity extracellular fluids (volume fraction of 15% to 30% [12]). In the study by Qi et al., uniform E-fields were imposed on the tissue specimen without considering the boundary charges between the specimen and surrounding bulk tissue, and therefore, the E-field required amplitude correction for proper comparison with the conventional homogenous method.

More fundamentally, the exogenous E-field generated by stimulation in brain tissue is not a directly measurable quantity. Therefore, using E-field as the source for comparison comes with a significant caveat. Only external sources, such as electrode voltages and currents or coil currents, are directly accessible. Experimental measurements and computational simulation [13] may report stimulation strength in terms of the total E-field that is calculated for a given stimulator output, but this E-field is a macroscopic equivalent and not the microscopic field in the sub-μm extracellular space (with typical widths of 40–80 nm [12]). Indeed, experimentally E-fields are calculated from the potential difference measured between pairs of microelectrodes, which are significantly larger than the extracellular space between cells.

Overall, this study highlights the importance of differentiating conductivity values obtained from the literature, depending on whether the space surrounding a neuron model is considered as bulk neural tissue (for studies with the two-step approach, e.g., [6]), or as the realistic micro-environment consisting of extracellular fluids (e.g., [9,10]).

### Matching field strength and conductivity

#### Methods

Qi et al. used brute force optimization to find the correction factor for the average extracellular E-field and to match the conductivity (section 2.8 of [1]). This was performed on an artificially generated rectangular cuboid sample due to the shape of the realistic tissue specimen being a non-orthogonal parallelepiped.

Previously, conductivity of retinal tissue was estimated by modeling the tissue layers between two parallel planar electrodes and solving the electrode voltages for current-controlled stimulation [14]. Least-square fitting was used to find the effective tissue conductivity in peripheral nerves [9], with the advantage of being applicable to any arbitrary and irregular geometry. In this method, the homogenous model is first solved with the same conductivity of the extracellular fluid *σ*_ex_ as in the microscopic tissue model. Then, the effective conductivity *σ*_eff_ of the homogenous tissue is obtained by scaling the solution to match that of the microscopic tissue model in the extracellular space using least-square fitting. For current-controlled stimulation, the fitting is conducted on the potentials *φ* [9]

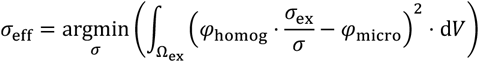

or E-field ***E***

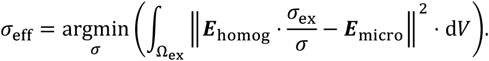

Here, the extracellular domain is Ω_ex_ and d*V* is the volume element for integration. For voltage-controlled stimulation, the current density could similarly be matched, whereas when the E-field is used as the source of stimulation as in the work by Qi et al., both the E-field and current density need to be matched.

Although the intracellular space is not involved in the above calculation, the results would not have changed if it is included, because after initial polarization [7], the intracellular E-field is zero and the intracellular potential equals the extracellular potential averaged over the enclosed cell membrane, whether the membrane is treated as an ideal insulator or not [9,10,14].

#### Models

To demonstrate the least-square matching method for a non-uniform E-field, we simulated the E-field distributions of spherical electrodes. The models were built in COMSOL (version 6.2, COMSOL, Inc.) and solved using the stationary electric current solver of the AC/DC module. Data analysis and additional visualization were performed in MATLAB (version 2024b, The MathWorks, Inc.). Key MATLAB code is provided in the supplement.

A spherical electrode with a radius of 50 μm was located at the origin, and the surrounding simulation space had a radius of 0.5 mm (Fig. 2A). Given the symmetry, only one octant of the space was represented, and the internal space of the electrode was removed to reduce computational costs. To reflect the symmetry, the artificial boundaries in the *x*–*y, x*–*z*, and *y*–*z* coordinate planes were set to insulation condition. The volume had a conductivity of 1 S/m and was grounded at the outer boundary. For current-controlled stimulation, the electrode surface was set to a floating potential with a current of 1 μA (i.e., 8 μA total for complete electrode), whereas for voltage-controlled stimulation, a fixed voltage of 1 V was applied.

**Figure 2.**
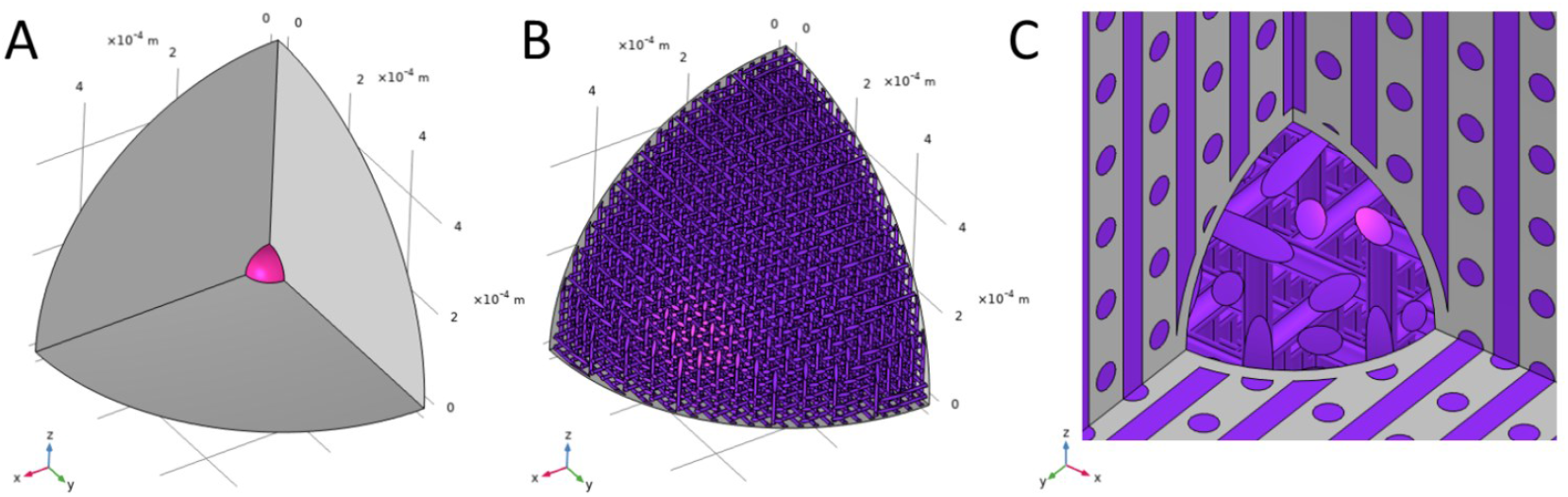
**A**. Homogenous tissue model with a spherical electrode (magenta) located at the center of the spherical simulation space, with only one octant explicitly modeled. The artificial boundaries in the *x*–*y, x*–*z*, and *y*–*z* coordinate planes (gray) are insulating. **B**. Microscopic tissue model consisting of three interleaved grids of parallel axons (purple). The axons do not intersect with the outer boundary. **C**. The axons closest to the coordinate axes are modified to prevent intersection with the electrode (not shown in this view).

For the microscopic tissue model, cylindrical axons were placed in three orientations parallel to the coordinate axes (Fig. 2B). For each orientation, the axons were arranged in a square grid with 25 μm center-to-center distance. The radius of the axons was 4.46 μm to achieve a volume fraction of 30%. To prevent the axons from intersecting with the spherical outer and electrode boundaries, they were truncated at an outer bound of 0.495 mm radius (Fig. 2B) and an inner bound of 55 μm radius (Fig. 2C), respectively. The intracellular and extracellular space were assigned the same conductivity of 1 S/m, and the membrane’s specific resistance was set to 1 Ω⋅m^2^ (i.e., specific conductivity 0.1 mS⋅cm^−2^, corresponding to 10 nS/m for a membrane thickness of 10 nm). Axon boundaries due to intersection with the coordinate planes were set to insulation boundary condition.

#### Results

For current-controlled stimulation, the electrode voltage for the homogenous and microscopic tissue models were 18.78 mV and 11.45 mV, respectively, indicating that the former should have a 39% lower conductivity of 0.610 S/m to match the latter. This was close to or slightly higher than the theoretical values for the 30% volumetric fraction between 0.552 and 0.609 S/m (range given for different formulas) [15,16]. Matching the solution globally provided similar results. The potential and E-field distributions of the two tissue models on the coordinate planes are shown in Fig. 3. Performing least-square fitting on the potential and E-field in the extracellular space resulted in an effective isotropic conductivity *σ*_eff_ of 0.602 and 0.642 S/m, respectively. For voltage-controlled stimulation, the total electrode current for the homogenous and microscopic tissue models were 87.26 μA and 53.20 μA, respectively. The homogenous tissue therefore should have a conductivity of 0.610 S/m, agreeing with the results for the current-controlled case. Least-square fitting on the current density in the extracellular space resulted in an effective isotropic conductivity *σ*_eff_ of 0.622 S/m.

**Figure 3.**
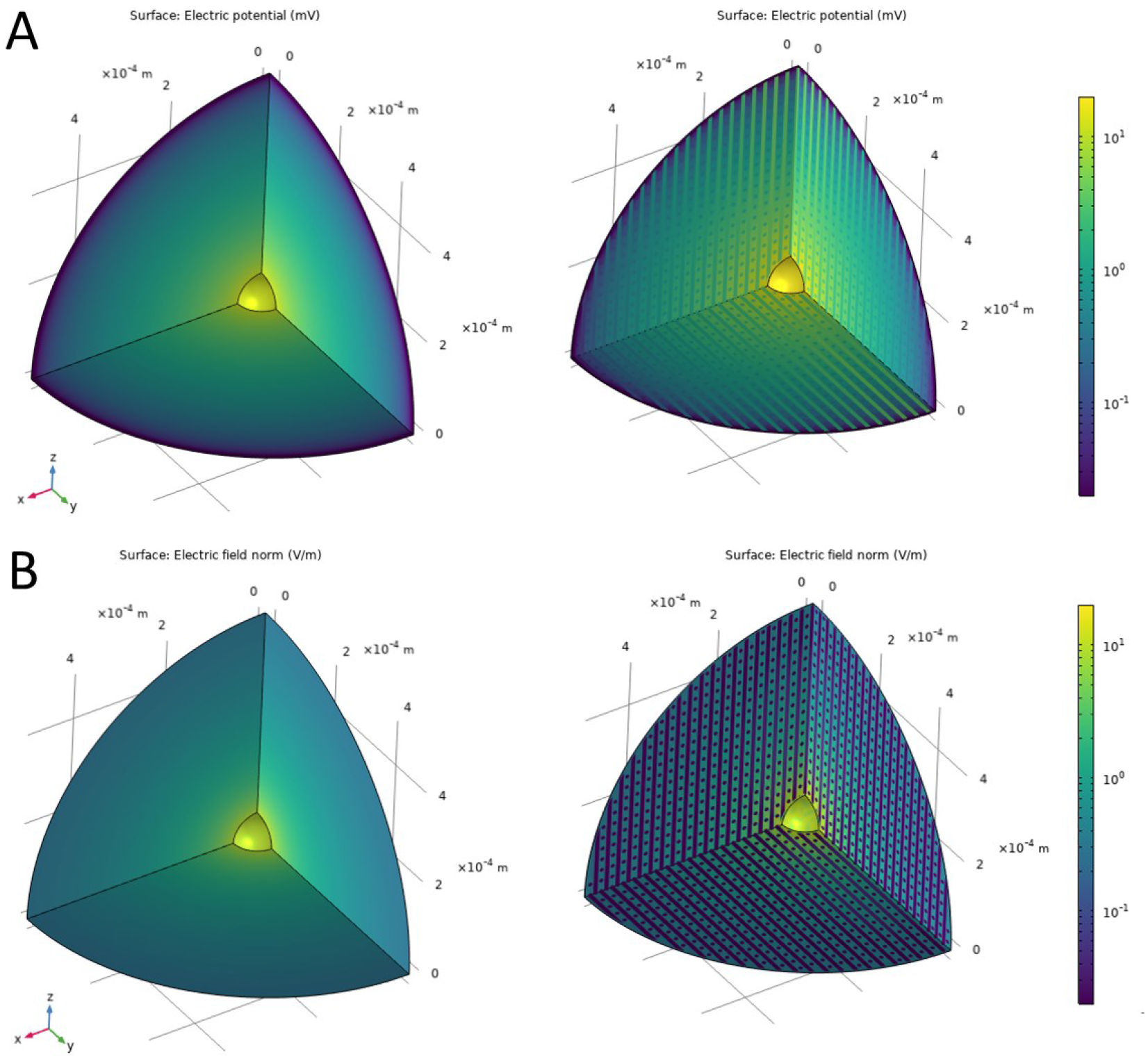
Distributions of the (**A**) electric potential and (**B**) E-field amplitude on the electrode surface and the coordinate planes of the homogenous model (left) and microscopic tissue model (right) for current-controlled stimulation before matching. Color maps are on logarithmic scales. The intracellular space of each axon has a uniform potential that equals the average extracellular potential on the membrane and the intracellular E-field is approximately zero.

Least-square fitting in the entire space resulted in similar conductivity values (Table 1). Performing the fitting with the potential was overall more numerically robust and accurate as well as more computationally efficient as the potential is a continuous scalar field, compared to the vector E-field and current density with large discontinuities across the membranes. A significant source of numeric inaccuracy came from the electrode’s vicinity, where the E-field was highly non-uniform, and the additional space to prevent intersection (with thickness of 10% electrode radius) resulted in the microscopic tissue not being homogenizable there.

**Table 1.** Calculation of equivalent conductivity using various methods.

| Equivalent conductivity of homogenous model (S/m) | Matching electrode voltage or current | Least-square fitting |  |
| --- | --- | --- | --- |
|  |  | Extracellular space | Entire space |
| Current-controlled | 0.6102 ($V$) | 0.6024 ($\varphi$)<br>0.6418 ($E$) | 0.6173 ($\varphi$)<br>0.6246 ($E$) |
| Voltage-controlled | 0.6098 ($I$) | 0.6422 ($J$) | 0.6010 ($J$) |

The potential, E-field, and activating function along a line parallel to the *x*-axis at *y* = 118 μm and *z* = 81 μm are shown for a cathodic current of −1 μA (Fig. 4). After matching the conductivity for the homogenous model (here using 0.6 S/m), the potentials agreed much better with the microscopic tissue. With the confounding factor removed, the remaining differences in the E-field and activating function were from the microscopic field perturbations, with their spatial extent and amplitude determined by the size of the axons that generated them. Although the peak amplitudes of the E-field and activating function were higher for the microscopic tissue model, the spatial noise averaged out to zero and resulted in the same amplitude profiles as those of the homogenous model. Given the spatiotemporal filtering of neural membranes, the threshold difference would therefore be reduced compared to the uncorrected case, i.e., the threshold reduction factor would be higher and closer to 1.

**Figure 4.**
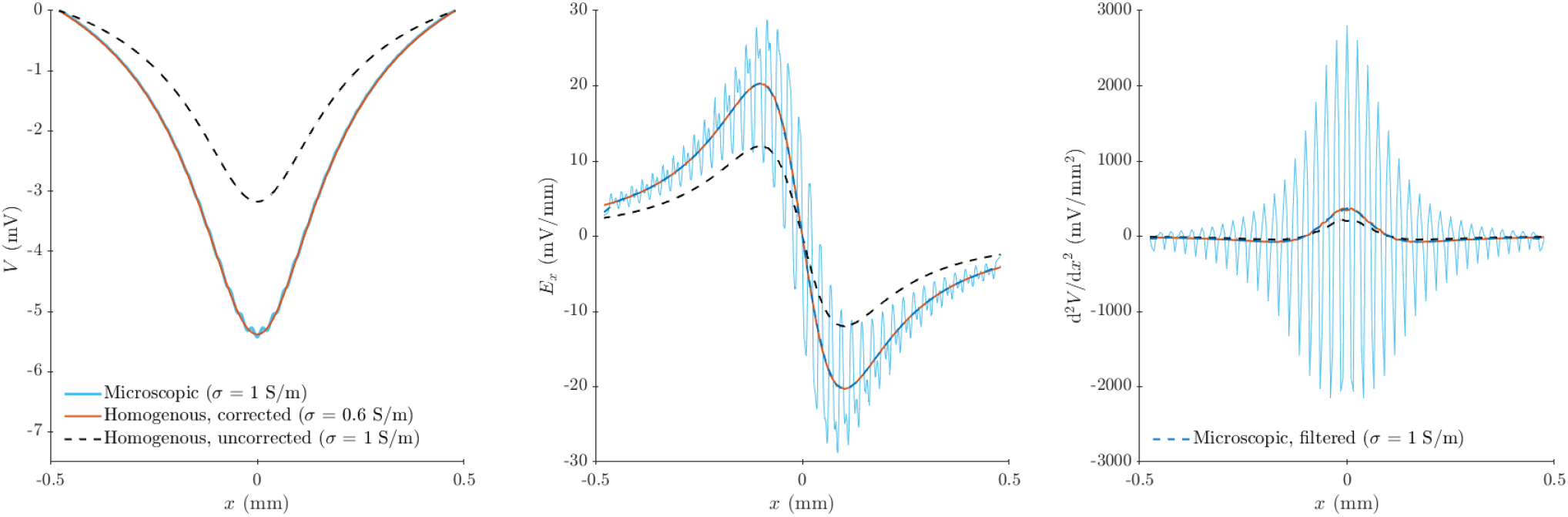
The potential (left), axial E-field (center), and activating function (right) along a line parallel to the *x*-axis at *y* = 118 μm and *z* = 81 μm. The data are extended (anti-)symmetrically to the negative *x*-axis for visualization. The potential and E-field are sampled from COMSOL with 1 μm resolution, and the activating function is calculated on the E-field in MATLAB as a finite difference with reduced resolution of 5 μm to suppress numeric noise. Compared to the microscopic tissue model (light blue lines), the uncorrected homogenous solution (black dashed lines) underestimates the strength of the stimulation significantly. After correcting for the conductivity, the solution of the homogenous model (orange lines) matches the average of the microscopic model. The dark blue dashed lines show the E-field and activating function filtered with a 4th-order Butterworth lowpass filter with cutoff spatial frequency of 10.6 mm^−1^, corresponding to a spatial constant of 15 μm that is much shorter than axonal length constants.

## Discussion and conclusions

Qi et al. further explored how the effect of E-field perturbations on activation thresholds changed with their spatial scales (section 4.3 of [1]). For a target axon with a length constant λ, the E-field perturbation to result in a noticeable effect on thresholds required large nearby neurites with diameters on the order of 0.1⋅λ. For realistic neurite sizes, the spatial scale of the perturbations was too small, and they diffused too quickly, to have an apparent effect. The spatiotemporal filtering of the membrane [17,18] also decreases the coupling of the microscopic E-field (Fig. 4). E-fields polarize neurites in proportion to the second spatial derivative of the collinear E-field, known as the *activating function*. It should be noted that for myelinated nerve fibers, which have the lowest thresholds due to stronger coupling with the E-field for a given fiber diameter [2,19], the effect of the microscopic noise on the coaxial E-field’s derivative is further diminished, because the activating function is effectively calculated as a second-order difference of the extracellular voltage over the internodal distance [6,19] of hundreds of micrometers to millimeters. Even for unmyelinated fibers, the microscopic E-field noise will potentially affect simulated neural responses only if the simulation uses spatial discretization that is smaller or comparable to the scale of the noise. While the authors explored extremely fine spatial and temporal resolutions (supplement C1 of [1]), such resolutions are not typically used in modeling studies, which may be justified based on these results.

The correction on the threshold reduction factors was only performed for one volume fraction value of 50% (section 2.8 of [1]). Applying the correction to all data points, the threshold reduction factors as a function of cell volume fraction (Fig. 8 of [1]) would likely be relative flat lines close to 1. If there were significant deviations from 1 in the mid-range (e.g., 30%–70% volume fraction), the function would certainly be nonlinear: at both 0% and 100%, the threshold reduction factors should equal 1.

The results by Qi et al. overall validated the conventional two-step approach for predicting activation thresholds: the thresholds on average did not change much due to the microscopic E-field perturbation. Although the local E-field could significantly increase in amplitude, whether such distribution outlier [20] would translate to significant decrease in thresholds still need to be validated, given that the high spatial frequency of the microscopic field perturbation result in much weaker coupling to the axons compared to locally uniform field.. Most computational models of cortical stimulation, which used the two-step approach, predict higher activation threshold than *in vivo* measurements [2,21]; this result suggests microscopic E-field perturbations are not the cause of this discrepancy, and instead, that other factors warrant investigation, including the lack of ongoing activity or inaccuracies in the membrane dynamics in the neural simulations. This is in agreement with recent work using a bidomain BEM method to simulate both the microscopic E-field perturbations and neural membrane dynamics in a single computational framework [22].

Recent advances in imaging and computation enabled dense volumetric reconstructions of brain tissue down to the nanoscale, which Qi et al. leveraged to simulate brain stimulation with unprecedented detail. While capturing this level of detail may not affect predictions of activation thresholds with the brief, uniform E-field pulses they tested, there are several exciting directions for this work moving forward. For example, the computational framework by Qi et al. also enables the numerical estimation of conductivity (potentially anisotropic) of macroscopic homogenous brain tissue from microscopically realistic brain samples. Further development could generate depth/layer specific neural tissue conductivities [14], for example using the larger cubic millimeter IARPA Phase III sample from mouse visual cortex [23] or the petavoxel human cortical sample [24]. This would then enable future FEM/BEM E-field simulation of brain stimulation to use macroscopically inhomogeneous conductivity and improve the solution accuracy. Additionally, the IARPA Phase III sample was imaged using two photon calcium imaging, combining nanoscale structural data and *in vivo* functional recordings from tens of thousands of neurons [23]. These multimodal datasets can not only reveal fundamental principles of cortical function but also serve as inputs to data-driven neuron and network models for addressing fundamental mechanistic questions of brain stimulation techniques. How are the diverse neuronal and non-neuronal cortical cell types affected by different stimulation paradigms or doses? How do their responses depend on their morphology or connectivity? How do the various cellular elements interact to generate macroscopic physiological responses (e.g., EEG or fMRI signals)? Building models to address these questions will require development and rigorous application of new computational frameworks. The study by Qi et al. represents one of the first steps in this direction, setting the stage for further advances in our understanding of brain stimulation.

## Acknowledgments

This work was supported by the US National Institutes of Health (NIH Grant No. R01 MH128422). The content is solely the responsibility of the authors and does not necessarily represent the official views of the funding agency. The authors declare no relevant conflict of interest.

## CRediT authorship contribution statement

B. Wang: Conceptualization, Investigation, Methodology, Formal analysis, Visualization, Writing – original draft, Writing – review & editing. A. S. Aberra: Conceptualization, Writing – review & editing.

## Supplement

The key MATLAB code for performing least-square fitting.

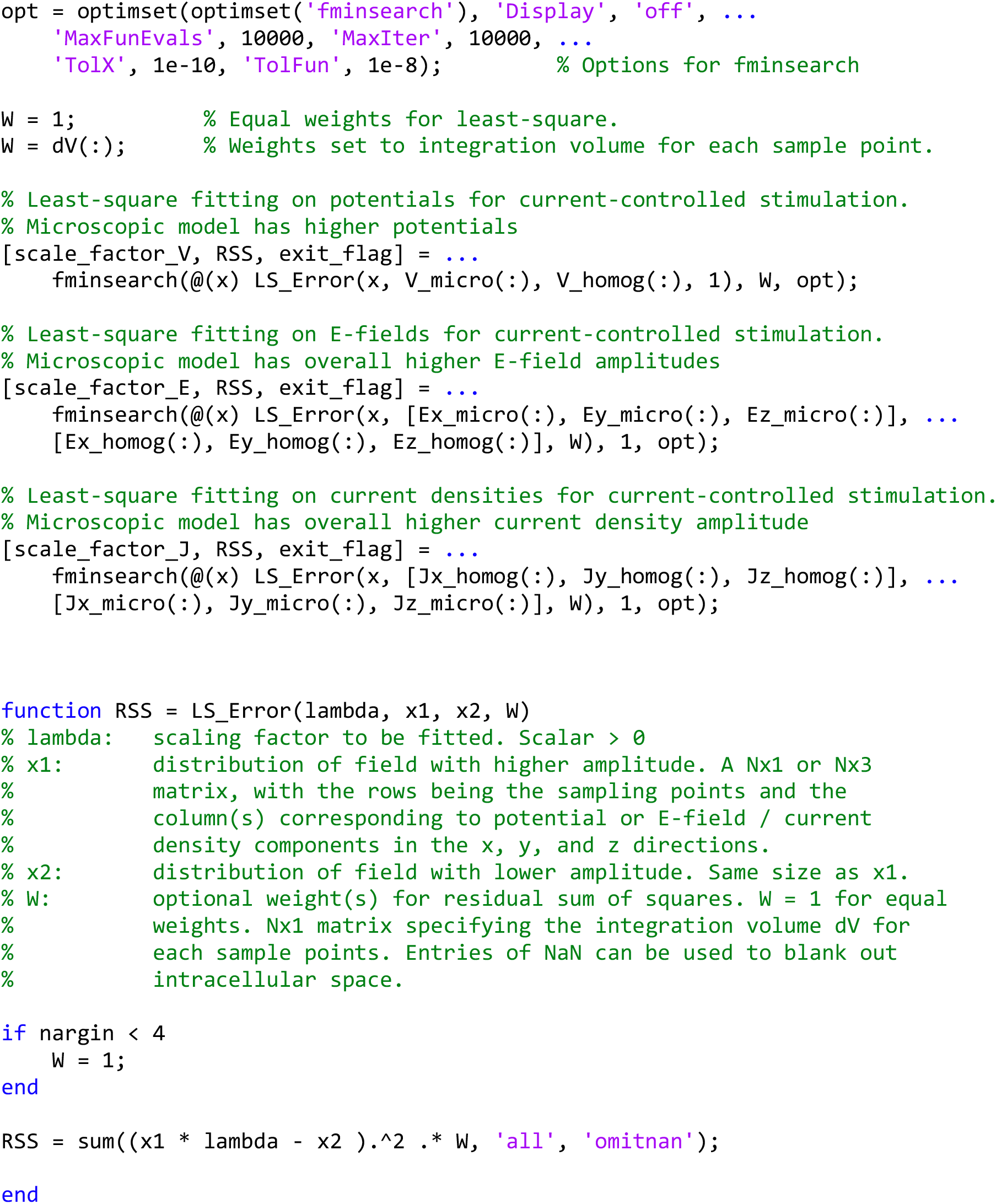

